# Structural insights into the insecticidal Vip3A toxin of *Bacillus thuringiensis*

**DOI:** 10.1101/2020.01.24.918433

**Authors:** Kun Jiang, Yan Zhang, Zhe Chen, Dalei Wu, Jun Cai, Xiang Gao

## Abstract

The vegetative insecticidal proteins (Vips) secreted by *Bacillus thuringiensis* are regarded as the new generation of insecticidal toxins because they have different insecticidal properties compared with commonly applied insecticidal crystal proteins (Cry toxins). Vip3A toxin, representing the vast majority of Vips, has been used commercially in transgenic crops and bio-insecticides. However, the lack of both structural information of Vip3A and a clear understanding of its insecticidal mechanism at the molecular level, limits its further development and broader application. Here we present the first crystal structure of the Vip3A toxin in an activated form. Since all members of this insecticidal protein family are highly conserved, the structure of Vip3A provides unique insight into the general domain architecture and protein fold of the Vip3 family of insecticidal toxins. Our structural analysis reveals a four-domain organization, featuring a potential membrane insertion region, a receptor binding domain, and two glycan binding domains of activated Vip3A. We further identify the specific glycan moieties recognized by Vip3A through a glycan array screen. Taken together, these findings provide insights into the mode of action of Vip3 family of insecticidal toxins, and will boost the development of Vip3 into more efficient bio-insecticides.

## Introduction

The entomopathogenic bacteria *Bacillus thuringiensis* (Bt), is the most widely used microbial insecticide in the world^1, 2^. It is renowned for its ability to produce insecticidal crystal proteins (Cry toxins) during its sporulation phase, which have been widely used in the prevention and control of agricultural pests through the development of transgenic plants or Bt-based biopesticides^3–5^. However, many pests are not sensitive to Cry toxins, and the development of insect resistance to Cry toxins has also been reported^1, 6–8^. The successful application of the Cry proteins, coupled with their limitations, has spurred on intensive research seeking to identify and characterize novel classes of insecticidal toxins that can be developed for agricultural purposes.

Vegetative insecticidal proteins (Vips), which are produced by Bt during its vegetative stages, have a wide spectrum of insecticidal activity, especially against lepidopteran pests^9^. To date, ∼150 distinct Vip toxins have been identified, which have been classified into four families (Vip1, Vip2, Vip3 and Vip4) based on their sequence similarity^10^. Among the Vip toxin family, Vip3A toxins are the most abundant and most studied^9^. Compared with known Cry toxins, Vip3A toxins share no sequence homology, bind to different receptors^11–14^, and lack cross-resistance^15–18^, therefore they are considered as ideal options to complement and expand the use of Bt in crop protection and resistance management. At present, the Vip3Aa toxin is the only family member that has been used in commercial transgenic crops together with Cry toxins, and no field-evolved resistance has yet been reported^1, 9, 19^. However, the lack of structural information and incomplete understanding of its mechanisms of action have severely limited the further development of Vip3A as a tool in pest control.

Vip3A toxins are large proteins (∼800 amino acids) consisting of a conserved N-terminus and a variable C-terminal region. The ∼89kDa Vip3A protoxin could be digested by insect midgut juices into two fragments: a ∼20 kDa fragment corresponding to the N-terminal 198 amino acids, and a ∼66 kDa fragment corresponding to the rest of Vip3A protein and carrying the toxic activity, which is considered to be the toxic core^9, 11, 20–25^. This processing step is regarded as an essential step for its activation and toxicity^11, 20, 21, 26–28^. Since their discovery in 1996^29^, Vip3A proteins have been the subject of intensive research. It has been reported that Vip3A stimulates membrane pore formation and apoptosis upon binding to target cells, which is proposed to be responsible for its cytotoxic effects^11, 30, 31^. The scavenger receptor class C like protein (Sf-SR-C) and the fibroblast growth factor receptor (Fgfr) have been reported as potential receptors for Vip3A^13, 14^. However, although some *in silico* modeling efforts and low resolution cryo-EM structures have attempted to obtain structural insight for these toxins, the atomic structure of Vip3A is still not available, which makes it difficult to reveal the relationship between their structure and function^27, 28, 32, 33^.

Here, we report the crystal structure of the toxic core of Vip3Aa. The structure shows a four-domain organization which is likely to be conserved for the toxic core of all insecticidal Vip3 family toxins. We identify conserved hydrophobic α-helices in domain II, which we predict to be involved in the membrane insertion process that follows the activation of cell-bound toxin. Structure-guided cell binding assays reveal that domain III is central to host cell targeting and binding of Vip3A toxins. Structural analysis and glycan array studies indicate that Vip3A toxins can recognize specific glycan moieties through domains IV and V. Together, our structural and functional studies provide new insights into the molecular mechanisms underlying mode of action of insecticidal Vip3 toxins.

## Results

### Overall structure of Vip3Aa11_200-end_

We used a Vip3A toxin from Bt strain C9, which has been named Vip3Aa11 (GenBank accession No. AY489126.1) in this study. Full-length Vip3Aa11 consists of 789 amino acids, which has been demonstrated to be digested between residues K198 and D199 by insect midgut juice^20, 26, 27^. Of the two resulting fragments, the C-terminal fragment (D199 to end) is considered to be the toxic core of Vip3A.

We started our crystallization trial with both Vip3Aa11 protoxin and Vip3Aa11 toxic core. Using spare matrix crystallization screening, we only identified one condition that yielded needle-shaped crystals of Vip3Aa11 protoxin. However, the crystals diffracted to only ∼15 Å and could not be improved despite extensive effort. No crystals were observed for the Vip3Aa toxic core construct despite screening more than 1000 crystallization conditions. However, when we deleted the N-terminal amino acid (Asp199) from the Vip3Aa11 toxic core, we obtained the crystal of Vip3Aa11_200-end_, which diffracted to ∼6 Å. Through the addition of an N-terminal MBP (Maltose Bind Protein) tag, we were able to isolate crystals with improved diffraction.

The final structure of Vip3Aa11_200-end_ was refined to a 3.2 Å resolution with *R* and *R*_free_ values to 0.1980 and 0.2389, respectively (Supplementary Table1). The crystal belongs to the P2_1_ space group and four MBP-Vip3Aa11 molecules were found in one asymmetric unit (Supplementary Fig. 1). These four molecules form two dimers in different orientations. PISA^34^ determined that there was limited interaction between two dimers, indicating that their association was caused by the crystal packing (Supplementary Fig. 1). Notably, the two Vip3Aa11 molecules in the “dimer” showed moderate conformational variations, with a core root mean square deviation (r.m.s.d) of 1.234 Å among 468 Cα atoms (Supplementary Fig. 2). Superimposition of separate domains between the two molecules revealed better alignment for domains III, IV and V, but not for domain II (Supplementary Fig. 2), suggesting that domain II might potentially be involved in the conformational changes during the activation of Vip3A toxins. However, the result of gel filtration chromatography showed that MBP-Vip3Aa11_200-end_ mainly exists as a monomer in solution (Supplementary Fig. 3), which is consistent with observations from a previous study^21^. Therefore, we used the monomeric structure of MBP-Vip3Aa11_200-end_ for subsequent analysis.

Our structure shows that the Vip3Aa11 toxic core is comprised of four domains (Fig. 1a and b). Therefore, the protoxin can be divided into five domains, starting from N-terminus: domain I, 1-198; domain II, 199-327; domain III, 327-518; domain IV, 537-667; and domain V, 679-789 in Vip3Aa11 (Fig. 1a and Supplementary Fig. 4). The overall structure of the Vip3Aa11 toxic core resembles a lobster, wherein domains II and III form the body, and domains IV and V are the claws of the lobster (Fig. 1b and c). The connection between domain II and domain III is compact. However, domains III/IV and IV/V are connected by long and flexible loops, which indicates that the relative locations and orientations of these two domains could change under different biological circumstances. There are over 100 known proteins of the Vip3 family. Based on their high degree of sequence conservation and previous studies^9, 20, 26, 27^, they are very likely to share similar overall structures and domain compositions.

**Fig. 1.**
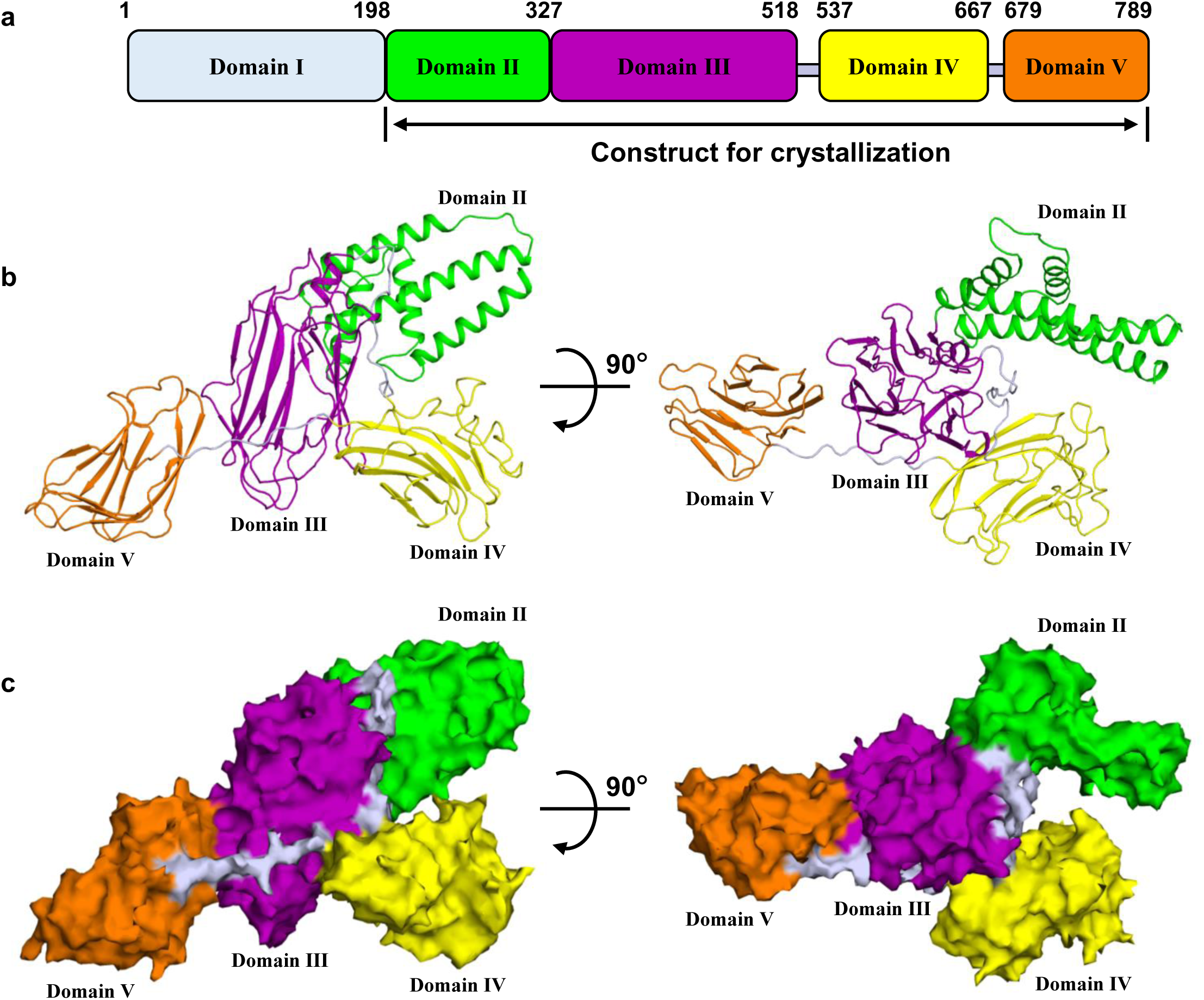
Overall structure of Vip3Aa11_200-end_. **a** Domain organization of Vip3A. **b** Two views of the overall structure of Vip3Aa11_200-end_ monomer coloured as in **a**. **c** Two views of the surface model of Vip3Aa11_200-end_ monomer coloured as in **a**.

### Domain II contains a conserved hydrophobic architecture reminiscent of a colicin A-type membrane insertion fold

Domain II of Vip3Aa11 (residues199-327) consists of five helices, which form two layers (Fig. 2a). The outer layer facing the solution contains two short helices, α2 and α3, while the inner layer that contacts domain III consists of three anti-parallel helices α1, α4 and α5. The outer layer contacts with the upper portion of the inner layer and is almost perpendicular to the inner layer. An intensive search against the PDB (Protein Data Bank) through the DALI server^35^ failed to identify any known structure that shows significant homology with all five α-helices of the domain II from Vip3Aa11, suggesting that this is a novel protein fold.

**Fig. 2.**
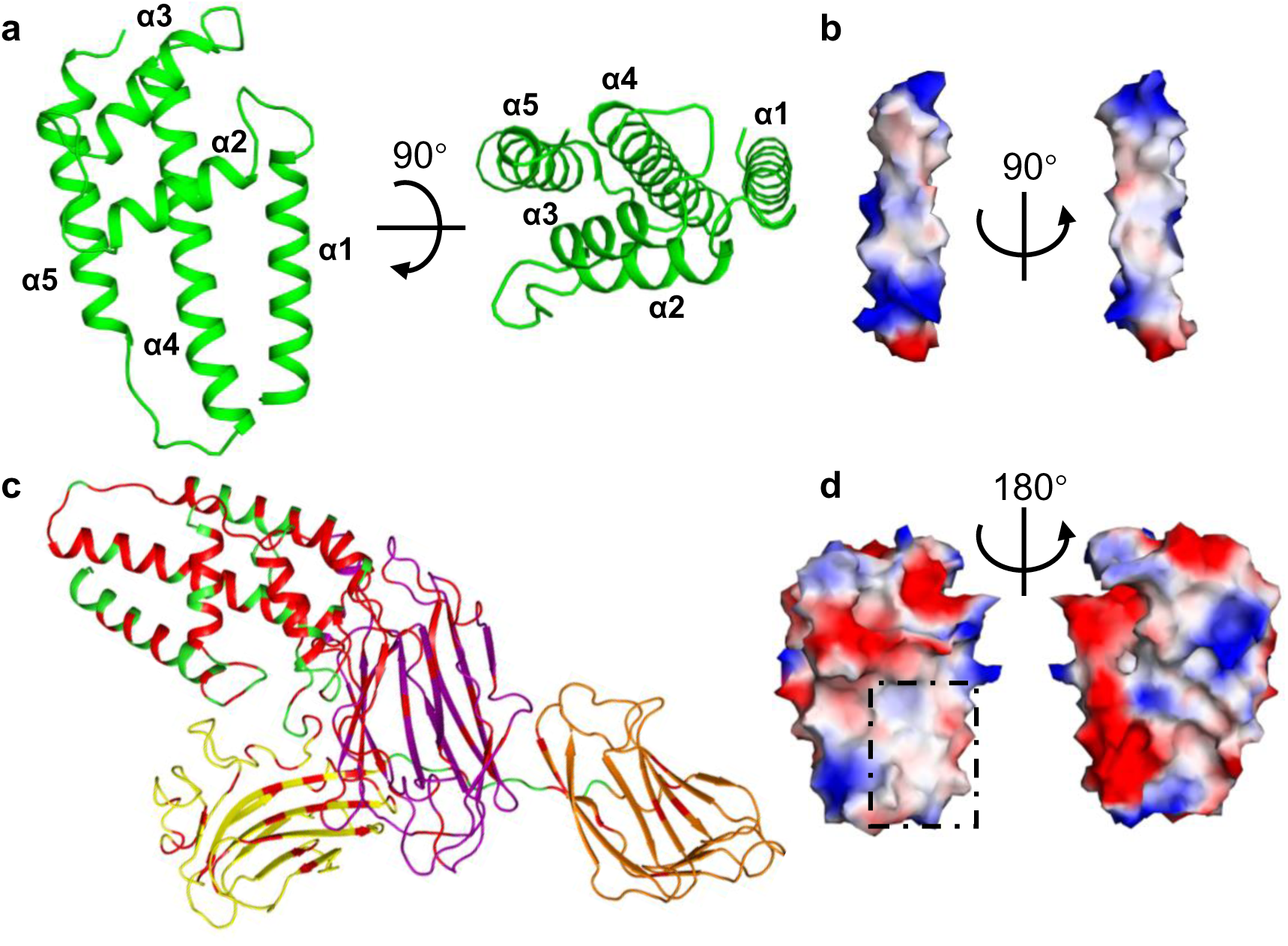
Domain II of Vip3Aa11 shows a conserved hydrophobic surface. **a** Two views of structure of Vip3Aa11 domain II shown as a ribbon cartoon. **b** Two views of the surface model of helix α4 from domain II show its surface charge distribution. **c** The highly conserved amino acid residues from Vip3 family sequence alignment (Supplementary Figure 4) are highlighted in the Vip3Aa toxic core structure with red color. **d** Two views of the surface model of Vip3Aa11 domain II show its surface charge distribution. The conserved hydrophobic surface is highlighted by black square. **b**, **d** The surface is coloured as the basis of electrostatic potential with positive charged surface in blue and negatively charged area in red.

Among these five helices, helix α4 is the longest one. It spans around 45 Å and contains 30 amino acid residues, starting from E267 at the N-terminus to L296 at the C-terminus. Electrostatic surface potentials analysis shows that the majority of charged and polar amino acid residues locate at N-terminal and C-terminal ends of helix α4 (Fig. 2b). For the middle portion of helix α4, from F274 to L289, 75% amino acid residues are hydrophobic residues. Sequence alignment through Vip3 family shows that the hydrophobic region of helix α4 is very much conserved and it is also the most agminated hydrophobic region of Vip3 family proteins (Supplementary Fig. 4). Close to helix α4, helix α1 also shows several conserved hydrophobic amino acid residues facing to helix α4 (Supplementary Fig. 4 and 5). Although these two α-helices are not directly connected in domain II of Vip3A, this structural feature is very similar to the hydrophobic helical hairpin from typical colicin fold, which has been shown to be involved in membrane insertion of several α-pore-forming toxins^36^.

Based on the sequence alignment, all the conserved amino acid residues were highlighted on the Vip3Aa11 toxic core structure (Fig. 2c). It also shows that domain II is the most conserved domain compared to other domains. Electrostatic surface potentials analysis shows that there is an obvious hydrophobic surface, which is mainly contributed by the conserved helix α1 and α4 (Fig. 2d).

This similarity, the strong conservation of this hydrophobic patch and previous data demonstrating the membrane insertion properties of Vip3A toxins^13^ collectively suggest that domain II is involved in the membrane insertion processes thought to be triggered once Vip3A binding to host cell receptors.

### Domain III is involved in receptor binding of Vip3A toxin

Domain III of Vip3Aa11 (residues 328-518) consists of twelve β strands and one short α-helix at the C-terminal end (Fig. 3a). Twelve β strands comprise three antiparallel β sheets sharing a similar “Greek-key” topology (Fig. 3a) with a hydrophobic center featuring highly conserved residues V349, F360, I362 and L370 from β sheet I, I425 and F427 from β sheet II and I481, F492 and L505 from β sheet III (Fig. 3b). The results from DALI server^35^ showed that the fold of domain III is similar to that of domain II of the Cry family of insecticidal toxins, which has been shown to be involved in host cell receptor recognition and binding^4^. We therefore sought to explore whether domain III serves as a receptor binding domain for Vip3A toxins.

**Fig. 3.**
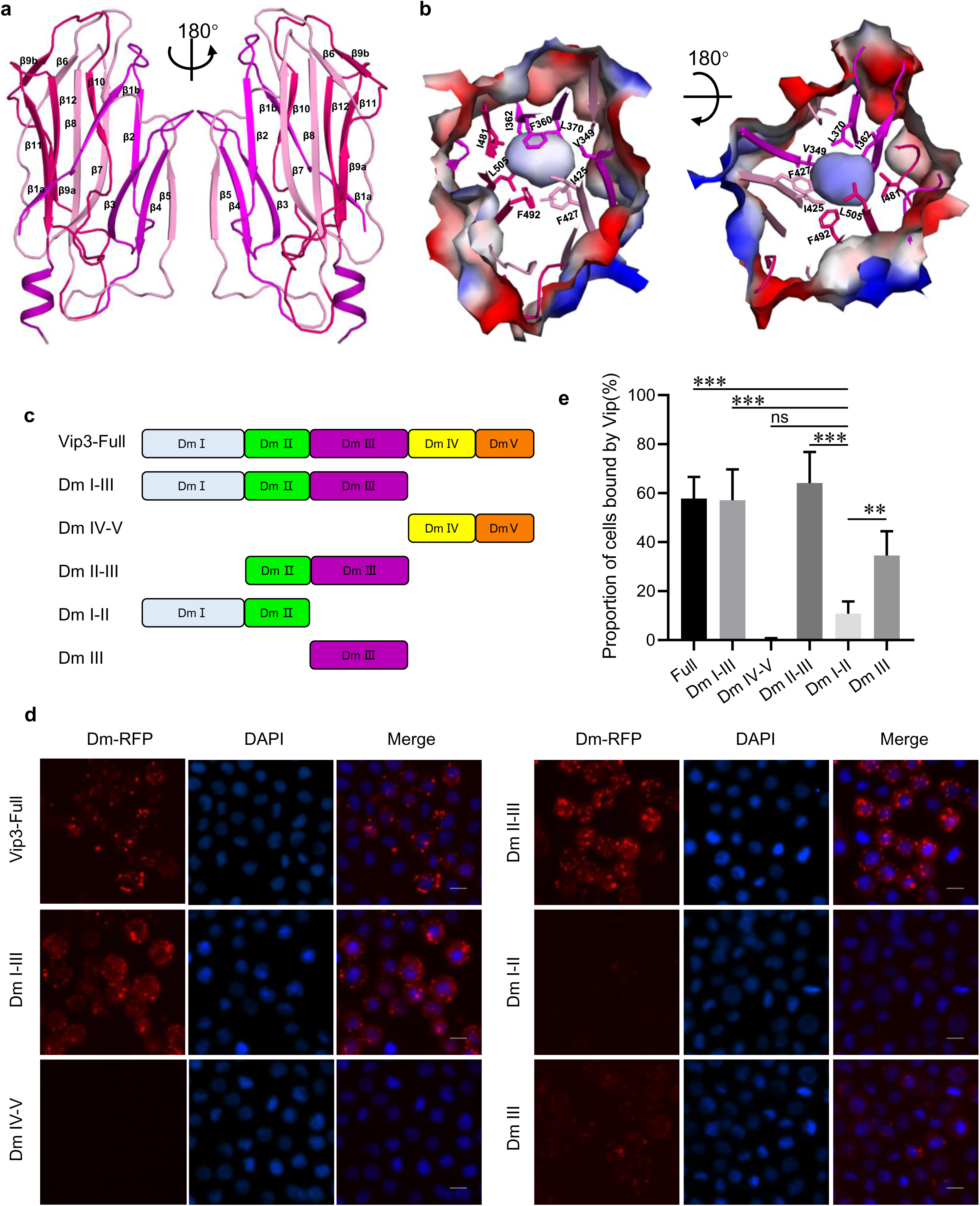
Domain III is a potential receptor binding domain. **a** Overall structure of Vip3Aa11 domain III shown as a ribbon cartoon. Two views of three antiparallel β sheets from domain III are shown in three different colors. **b** Two views of the surface model of domain III of Vip3Aa11. Inside the domain III, there is a conserved hydrophobic core, and the conserved hydrophobic amino acid residues from three antiparallel β sheets are shown as sticks. **c** The schematics of C-terminal RFP tagged of Vip3Aa and its truncation derivatives. **d** Fluorescence microscope images of Sf9 cells treated with Vip3Aa-RFP or its truncations, which were labeled with C-terminal RFP tag, for 6 h. Nuclei are stained with DAPI (blue). **e** Quantification of the number of Sf9 cells that can be bound by RFP-tagged Vip3Aa and its truncations of Fig3**d** in a blind fashion (n = 100 cells per sample). Data are expressed as the mean ± SD from three independent experiments.

To explore this hypothesis, we used *Spodoptera frugiperda* cells (Sf9 cells and Sf21 cells), which have previously shown to be specifically targeted by Vip3 toxins^14, 37^. To determine which domain(s) of Vip3Aa11 interact with Sf9 cells, we carried out fluorescence-based cell binding assays using different C-terminal RFP-tagged Vip3Aa truncation derivatives (shown schematically in Fig. 3c). As shown in Fig 3d, 3e and Supplementary Fig. 6, while domains IV and V do not show detectable binding to Sf9 cells, the binding of a construct featuring only domains II and III to Sf9 cells is indistinguishable from that of full-length Vip3Aa. The interaction of domain III alone with Sf9 cells is significantly stronger than that of the domain I-II construct, indicating that domain III is central to Vip3A receptor binding to Sf9 cells.

### Domains IV and V are glycan binding motifs

Domains IV and V both are all β sheets folds (Fig. 4a and b). Unlike domains II and III, which have compact organization, domains III/IV and IV/V are connected by long and flexible loops (Fig. 4a). In addition to these loops, there are several polar interactions between domains IV/V and domain III, that reduce the flexibility and fix the domains IV and V at the observed positions and orientations (Fig. 4a).

**Fig. 4.**
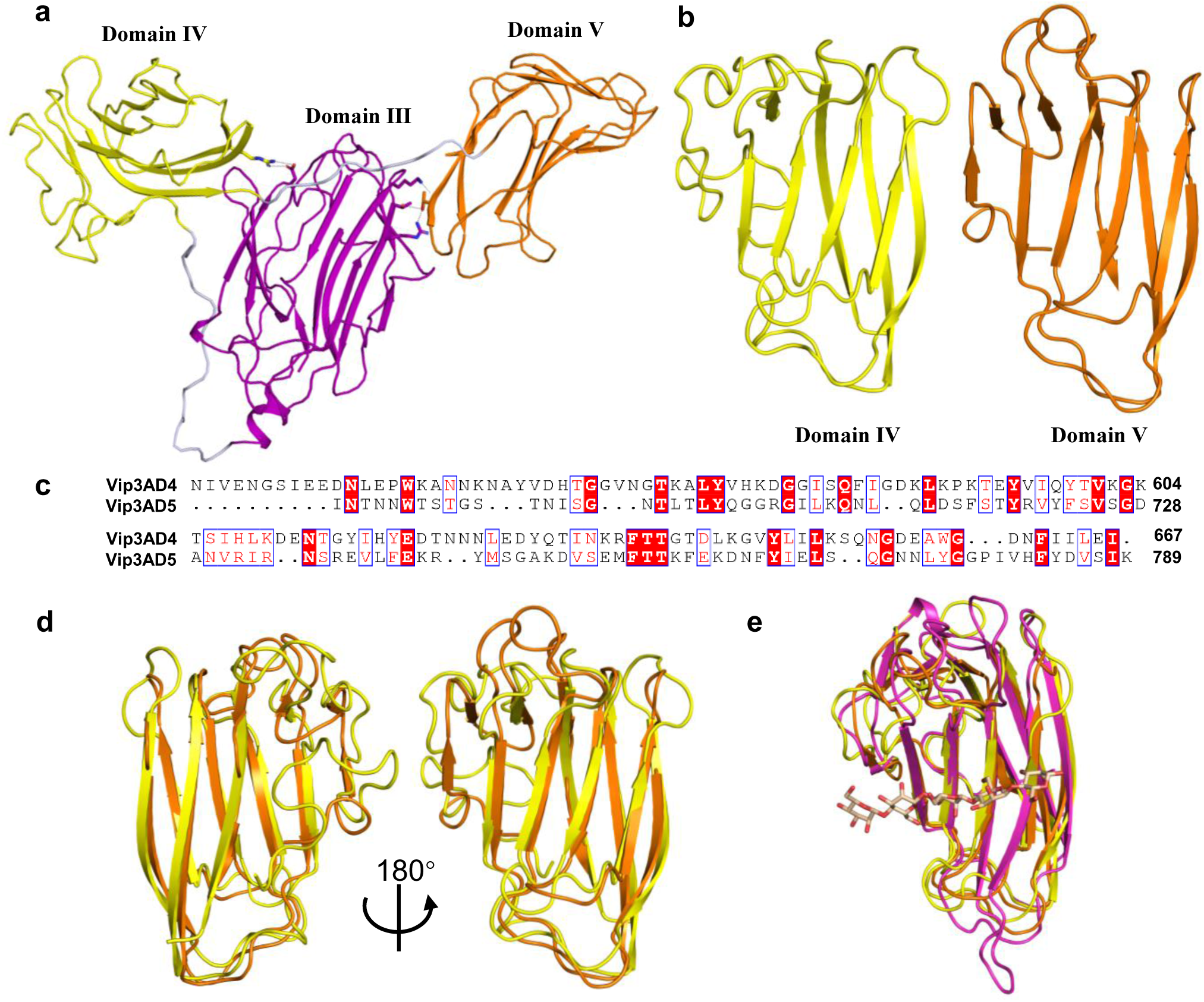
Domains IV and V of Vip3Aa11 have glycan binding motifs. **a** Domain architectures of domains III, IV and V of Vip3Aa11. The polar interactions between domain IV, V and domain III are shown as sticks. **b** Overall structure of Vip3Aa11 domain IV and V shown as a ribbon cartoon. **c** Amino acid sequence alignment between domain IV and domain V of Vip3Aa11. The highly conserved amino acid residues are shaded in red color. ClustalX2 was used to do the sequence alignment^57^. ESPript-3.0 was used to generate the figure^58^. **d** Two views of structure superimposition between domain IV and domain V of Vip3Aa11 shown as a ribbon cartoon. Color of each domain is consistent with Figure4**b**. **e** Structure superimposition between domains IV, V of Vip3Aa11 and glycan bound CBM16 (family 16 carbohydrate binding module, RCSB ID 2ZEY) shown as a ribbon cartoon. Domains IV and V are coloured as Figure4**b**, and CBM16 is shown in magenta color. The glycan in CBM16 is shown as stick in light brown color.

Domains IV and V are both built from two anti-parallel sheets of β sandwich, forming the “jelly-roll” topology. Despite showing only 17% sequence identity (Fig. 4c), domains IV and V align very well structurally, with a root-mean-squared deviation (r.m.s.d) of 1.299Å over 61 Cα atoms (Fig. 4d). To examine the potential function of these two domains, we searched for their structural homologues using the DALI sever^35^. The results for both domains show a very high similarity (Z score>10) to family 16 carbohydrate binding module (CBM16) of S-Layer associated multidomain endoglucanase (RCSB ID 2ZEY). Superimposition of domains IV, V and CBM16 demonstrates that these three motifs share a similar fold (Fig 4e), suggesting that they likely share a related function as well. CBM16 is a carbohydrate-binding domain of the highly active mannanase from the thermophile *Thermoanaerobacterium polysaccharolyticum* with high specificity toward β1,4-glucose or β1,4-mannose polymers^38^. Analysis of the electrostatic surface potential shows that both domains IV and V have a surface pocket at a similar position to a sugar-binding pocket of the CBM16 domain, although all three pockets have different shapes and charge distributions (Fig. 5a). Taken together, our structural analysis indicates that domains IV and V of Vip3A both contain a conserved glycan binding motif and that these motifs may target different sugars.

**Fig. 5.**
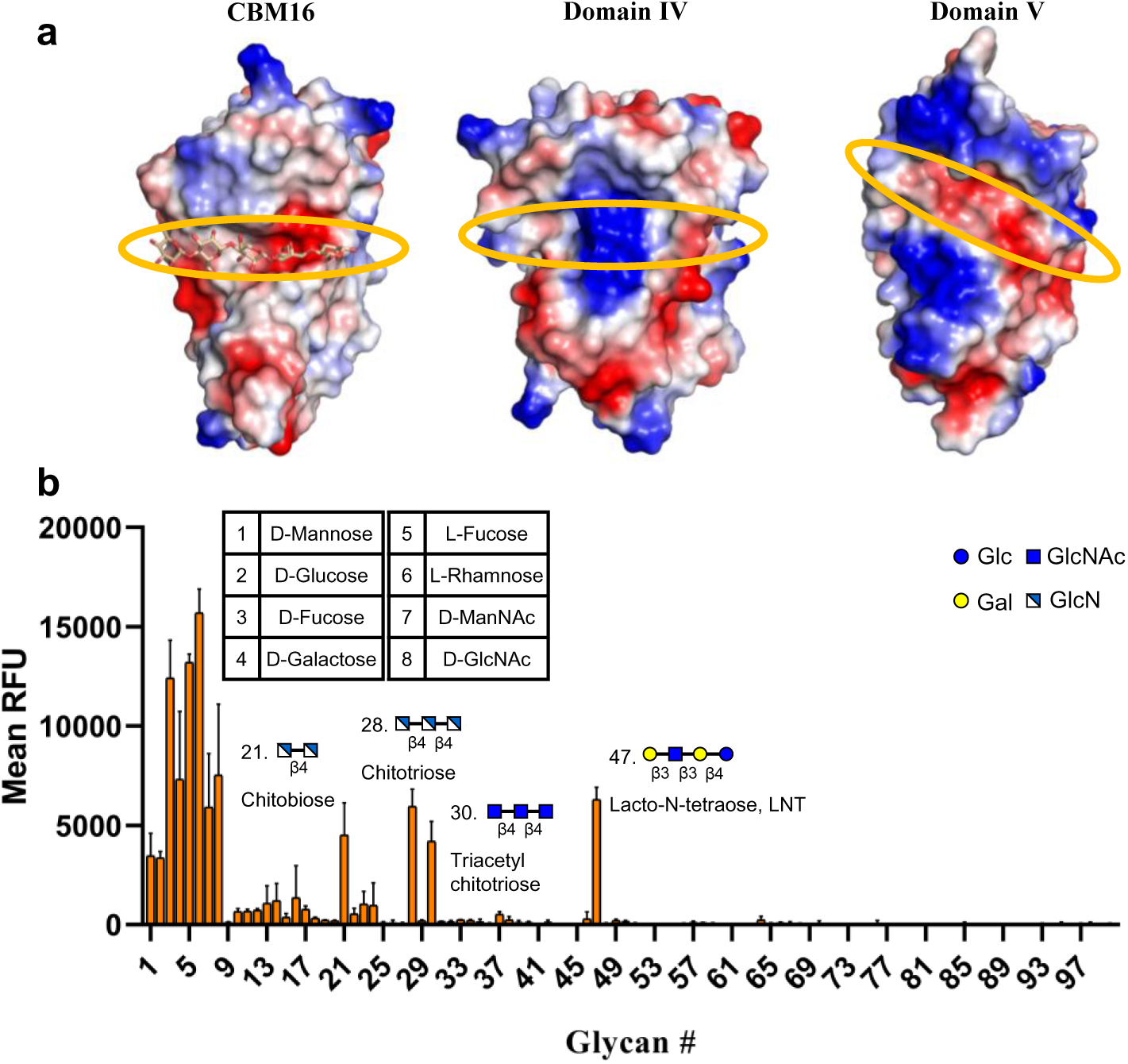
Domains IV and V of Vip3Aa11 can recognize glycans. **a** Surface charge distribution of the sugar-binding pocket of CBM16 and potential sugar-binding pocket of domains IV, V of Vip3Aa11, highlighted with the orange circle. **b** Glycan array analysis of Vip3Aa11 binding on a single glycan array 100 chip. Values represent the average relative fluorescence unit (RFU). The x axis depicts the glycan numbers. The structures of the most relevant glycans are shown.

In order to investigate the glycan binding properties of Vip3Aa11, we conducted glycan array screening with purified biotin-labelled, full-length Vip3Aa11. Screening results revealed that Vip3Aa11 has a strong binding preference to chitosan (GlcNβ1-4GlcN, Chitobiose and GlcNβ1-4GlcNβ1-4GlcN, Chitotriose) and chitin (GlcNAcβ1-4GlcNAcβ1-4GlcNAc, Triacetyl chitotriose) (Fig. 5b and Supplementary Table 2). Chitin is a major structural component of the peritrophic matrix (PM), an extracellular envelope that surrounds the midgut of most insects^39^. Epithelial cells in the midgut of insects synthesize chitin as a homopolymer of β-(1,4)-N-acetyl-D-glucosamine (GlcNAc), which may be subsequently converted to chitosan (a homopolymer of GlcN) by chitin deacetylase (CDA) secreted by insect midgut cells^40^. The identification of chitin and chitosan as targets of Vip3Aa is therefore consistent with previous studies that this toxin demonstrates specificity for binding insect midgut cells ^25, 41^. It also gives a potential explanation that why domains IV and V could not bind to the sf9 cells which are from ovarian tissue and may lack of chitin and chitosan^39, 40^. In addition to chitin and chitosan, the glycan array results revealed that Vip3Aa binds Lacto-N-tetraose as well as several monosaccharides including 6-deoxy-hexose, D-Fucose, L-Fucose and L-Rhamnose (Fig. 5b). Further studies will be required to determine the physiological relevance of Vip3A’s glycan binding specificities.

## Discussion

Vip3A toxins show a wide spectrum of specific insecticidal activities and are functionally distinct compared to the Cry toxins. These features makes them good candidates for combined application with Cry toxins in transgenic crops to broaden the insecticidal spectrum and to prevent or delay resistance^1, 9, 14^. The structural features and insecticidal mechanisms of Cry toxins have been studied in detail, which has been crucial to their widespread application^2–6, 19^. However, despite of the fact that Vip3 toxins were identified almost 25 years ago^29^, their mode of action remains poorly understood. One of the main reasons for this is the lack of a high-resolution three-dimensional structure, which significantly impedes detailed molecular level functional and mechanistic studies and thus limits the development of their insecticidal potential. Accordingly, many groups have made great efforts to obtain or predict the atomic structure of Vip3A^21, 27, 28, 32, 33^. However, high-resolution crystal structure of Vip3 toxin family is still missing. In this study, we report the first crystal structure of the Vip3A toxin, which provides a badly-needed framework to explore the molecular-level functional details of Vip3-family toxins.

Although the amino acid sequence similarity between the Vip3 family toxin and the three domain Cry family (3d-Cry) toxin is very low, our three-dimensional structural analysis showed interesting convergent evolution between these two families. Domain II of Vip3 has an all α-helix fold, including two conserved hydrophobic α-helices. Similarly, domain I of 3d-Cry also has an all α-helix fold and two hydrophobic α-helices, although it has additional α helices surrounding the conserved hydrophobic helices^4, 42^. Several studies have reported that domain I of 3d-Cry toxin is involved in its membrane insertion and pore formation processes through its conserved hydrophobic α-helices^4, 43–45^. This therefore suggests that domain II of Vip3 may also take part in these processes through its conserved hydrophobic α-helices. This is further supported by the fact that the insertion of the typical colicin fold, a hydrophobic helical hairpin, into the inner bacterial membrane is a minimal requirement for the pore formation of the pore-forming colicins^36^.

Both domain III of Vip3 and domain II of 3d-Cry are comprised of three β sheets with a conserved hydrophobic core. Extensive studies on domain II of 3d-Cry toxins showed that it plays a key role in the recognition of midgut receptors. The results of our cell binding assay indicate that Vip3 domain III is also central to cell binding. However, the receptor binding specificity of Vip3 toxins is still unclear and requires further studies.

Domain III of 3d-Cry toxins was predicted to bind glycans with a classic glycan binding motif^46–49^. Based on the amino acid sequence analysis, previous studies also predicted that all Vip3A proteins contain a carbohydrate-binding motif (CBM_4_9 superfamily; pfam02018) in the C-terminus (amino acids 536 to 652 in Vip3Aa)^9^. The function of this glycan binding domain is not yet clear^9^. In the present structure, we found that instead of the single CBM found in Cry toxins, there were two different CBM domains in the C-terminus of the Vip3 toxin, forming domains IV and V respectively. Our structural analysis indicates that the putative glycan-binding pockets of these two domains differ significantly, suggesting that they are likely to have different glycan binding specificities. This multiplicity of CBMs in Vip3 toxins may increase the diversity of their target polysaccharides. Interestingly, our glycan array results indicate that Vip3Aa is able to bind both chitin (and the related chitosan) as well as a spectrum of different monosaccharides. It is tempting to speculate that each of these two classes of glycans is specifically recognized by one of Vip3’s two glycan-binding domains.

In conclusion, we find here that although the overall structure and domain organization are very different between Vip3 toxin and 3d-Cry toxin, these two families are comprised of functionally and structurally related modules that are assembled in different ways, which may expand the insecticidal spectrum of Bt and make Bt more powerful and efficient to target and kill their hosts.

During the preparation of this manuscript, Zheng et al. reported the crystal structure of full-length Vip3B2160 protein^50^, which shares around 60% sequence identity to Vip3A. The overall structure of Vip3B2160 showed a five-domain organization (Fig. 6a). The domain I of Vip3B2160 (the pro-toxin domain cleaved to produce the active core that we have crystalized here) formed a unique fold containing five α-helices wrapping around Domain II (Fig. 6a). When these two Vip3 protein structures are superposed, domains III, IV and V of Vip3B2160 have very similar folds and organization with their counterparts from Vip3Aa11, respectively (Supplementary Fig. 7). However, there are dramatic differences in their domain II conformations, especially for the helices α1 and α2 (Fig. 6b). In Vip3B structure, similar to classic pore-forming toxin colicin A^36^, the highly conserved hydrophobic α-helix (corresponding to the helix α4 in Vip3A domain II) is surrounded by other helices from domains I and II. We hypothesize that this structural difference in domain II between the full-length and cleaved Vip3 proteins may represent the conformational change that switches this toxin from “inactive” to “active”. In this scenario, once domain I is cleaved by insect midgut juice, the α-helices of domain II undergo a dramatic structural shift that enables helix α1 to rotate to form a hairpin-like structure with helix α4. This new conformation, which strongly resembles the colicin A-type membrane insertion fold, represents the active membrane insertion moiety of Vip3 toxins (Fig. 6c). In a word, our crystal structure of Vip3A toxic core, combined with the structure of full-length Vip3B, reveals that the cleavage between domain I and domain II of Vip3 toxin, and the further conformational changes of domain II are essential steps for Vip3’s activation and function.

**Fig. 6.**
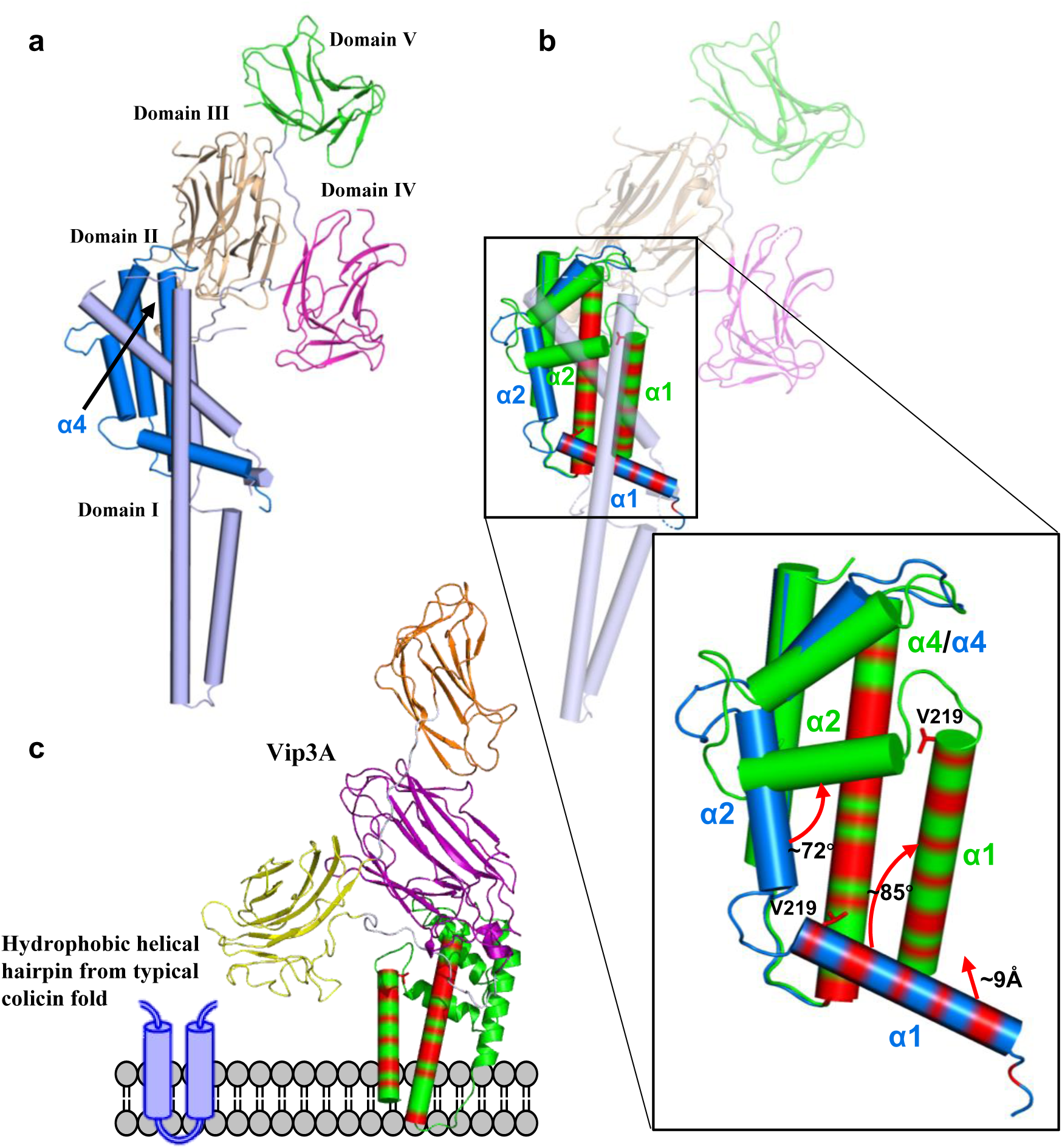
Proposed conformational changes of Vip3 toxin after the cleavage of domain I. **a** Overall structure of Vip3B2160. Domain I and II are shown as a cylindrical cartoon. Domain III, IV and V are shown as a ribbon cartoon. Domains I, II III, IV and V are coloured in light blue, blue, light brown, magenta and green, respectively. **b** Structural overlay of domain II between Vip3A(green) and Vip3B(blue). The inset shows a detailed view of the conformational changes. Appreciable conformational differences are observed in helices α1 and α2. The conserved hydrophobic amino acid residues are marked in red on helices α1 and α4. The Val219 is highlighted as stick to show the rotation along its y axis of helix α1 during the conformational change. **c** A proposed membrane insertion model of Vip3A toxin. A conserved hydrophobic helical hairpin from typical colicin fold is shown in blue (left-hand side). The conserved hydrophobic helices α1 and α4 of Vip3A form a hairpin-like motif and are shown as cylindrical cartoon. Similar to the colicin family pore-forming toxin, Vip3A may insert into the host cell membrane through the hairpin-like structure from domain II.

In summary, we report here the first crystal structure of Vip3A toxic core. It revealed the general domain organization of Vip3 family toxin and the potential function of each domain. Our study confirmed the glycan binding ability and defined the specificity of the Vip3A toxin using a glycan array screen. Importantly, we identify a conserved colicin A-type membrane insertion fold in the active core of our Vip3A structure which, through a structural comparison with full-length (inactive) Vip3B, enabled us to propose an activation mechanism for the Vip3 family toxins. Collectively, these data provide important structural and functional insights into Vip3 family toxins as well as a valuable resource to guide future studies and to re-evaluate the previous genetic and functional studies that will be crucial for the development of Vip3 as a new generation of bio-insecticides.

## Materials and methods

### Bacterial strains, cell lines and plasmids

*E. coli* BL21(DE3) for plasmid constructions and protein purification were cultured at 37 °C in lysogeny broth (LB) or agar. Methionine auxotrophic *E. coli* strain B834 (DE3) (Novagen) were used for selenomethionine-substituted (SeMet) Vip3Aa_200-end_ expressing. The *S*. *frugiperda* Sf9 cells were maintained and propagated in Sf-900 II SFM (Invitrogen) culture medium at 27 °C. The DNA of Vip3Aa_200-end_ was amplified from the Vip3Aa11 gene (GenBank accession No. AY489126.1) using oligonucleotide primer Vip200-F and Vip200-R and cloned into the pET28a vector with an N-terminal 6×His-MBP (maltose binding protein) tag. Plasmids used for RFP (red fluorescent protein) and C-terminal RFP tagged Vip3Aa (Vip3Aa-RFP) expression were constructed as described by Kun et al^14^. The different Vip3Aa truncations DNA were amplified from the Vip3Aa11 gene using oligonucleotide primer pairs; DmI-III-F and DmI-III-R, DmIV-V-F and DmIV-V-R, DmI-II-F and DmI-II-R, DmII-III-F and DmII-III-R, and DmIII-F and DmIII-R, and cloned into the pET28a vector with an C-terminal RFP-6×His tag, respectively. All plasmids were generated by the Gibson assembly strategy^51^. The nucleotide sequences of recombinant plasmid were verified by DNA sequencing. All the primers used in this study are shown in Supplementary Table 3.

### Protein expression and purification

Native His-MBP-Vip3Aa_200-end_ (Vip3Aa_200-end_) protein was expressed in *E. coli* B21(DE3) at 25°C for 48h in autoinduction Terrific broth (TB) medium. The cells were harvested by centrifugation at 5000×g at 4 °C for 15 min and the pellet was resuspended in lysis buffer (20 mM Tris–HCl pH 8 and 150 mM NaCl). After the cells were lysed by high pressure cell crusher (Union-Biotech co,.LTD), the supernatant was collected after centrifuged at 12,000×g at 4 °C for 60 min. The proteins were purified using Ni-NTA agarose resin (Qiagen), washed with 20 mM Tris-HCl, 150 mM NaCl, 20 mM imidazole, pH 8.0, and then eluted with 300 mM imidazole. The Vip3Aa_200-end_ proteins were further purified by HiTrap Q HP ion-exchange chromatography and Superdex 200 gel filtration chromatography (GE Healthcare Life Sciences). Fractions containing the Vip3Aa_200-end_ protein were concentrated to ∼ 7 mg/ml for crystallization. The expression and purification steps of other Vip3Aa truncations were the same as those of Vip3Aa_200-end_.

SeMet-substituted Vip3Aa_200-end_ was expressed in *E. coli* B834(DE3) strain. Briefly, the cells were cultured in the LB medium at 37 °C with shaking until the OD600 of the bacterial culture reached 1.0. The cells were harvested by centrifugation at 4000×g at 4 °C for 15 min and the pellet was washed one time with PBS. The pellet was resuspended in 1 L Medium A (M9 medium plus) and incubated for 3 hours at 37 °C. Added 50 mg seleno-methionine in the medium and incubated for a further 30 minutes. The protein was incubated to express for a further 10 hours by adding 200 mM IPTG (isopropyl-β-D-thiogalactopyranoside). The SeMet-Vip3Aa_200-end_ was purified by the same procedure as for the native Vip3Aa_200-end_ protein.

### Crystallization and data collection

The purification of His_6_-tagged MBP-Vip3Aa_200-end_ used for crystallization is described above. MBP-Vip3Aa_200-end_ (5 mg/mL) was used to perform initial spare matrix crystal screening with a crystallization robot. After crystal optimization trials, MBP-Vip3Aa_200-end_ (7 mg/mL) crystals grew in 3 days at 18℃ using the hanging-drop vapor-diffusion method in a mix of 1 μl of protein with 1 μl of reservoir solution consisting of 0.1 M sodium acetate pH 4.2, 0.5 M potassium formate, 0.1 M ammonium sulfate and 11% PEG4000. SeMet MBP-Vip3Aa_200-end_ crystals grew in the similar condition.

A native data set with the space group of P2_1_2_1_2_1_ was collected at 3.62 Å (native I). A weak selenomethionine (SeMet) derivative data set was collected at 3.9 Å with the same symmetry as the native I crystal for the amino-acid assignment using the difference Fourier map of the SeMet derivative. After further crystallization optimization, another native crystal (native II) was obtained with the space group of P2_1_ that could diffract to around 3.2 Å. Diffraction data were collected on BL17U1 and BL18U beamlines at Shanghai Synchrotron Radiation Facility (Shanghai, China) and processed by HKL2000^52^.

### Structural determination and refinement

Molecular replacement was carried out to identity the MBP positions in the native crystals by PHASER^53^. The initial phases were further improved with the multi-crystal averaging^54^. Model building was performed manually in COOT^55^, and the sequence assignment was helped with the SeMet anomalous difference map. Figures were prepared using PyMol (v.2.3.2, https://pymol.org/). Structure refinement was done by PHENIX^56^. The data collection and refinement statistics are summarized in Supplementary Table 1.

### Glycan microarray analysis

Purified full-length Vip3Aa protein were labelled with EZ-Link NHS-LC-LC-Biotin reagent in 20 mM HEPES pH 8.0 and 100 mM NaCl with a 4:1 molar excess of biotin. Unreacted biotin was removed by dialysis, and the resulting protein was concentrated by Amicon Ultra centrifugal filter (Millipore). The sample was diluted to 2.5 μg/ml and analyzed on a glycan 100 binding array provided by Creative Biochip. The reaction buffer was used as the negative control and the streptavidin was used as the positive control. The signals were measured in microarray scanner (LuxScan-10K/A; CapitalBio). The mean signal representing one glycan probe is the average of signals from triplicate spots on a single glycan array and compared by one-way ANOVA with Dunnett’s multiple comparisons test. All assays were repeated at least three times.

### Immunofluorescence

Sf9 cells with a density of 5 × 10^4^ cells per ml were seeded into 6-well culture plates separately. After overnight culture, the cells were respectively treated with RFP tagged Vip3Aa or its truncations (0.15 μM) for 6 h. After treatment, the cells were washed three times with PBS to remove unbound proteins, and fixed with 4% paraformaldehyde at 37 °C for 15 min. The cell nuclei were labeled with DAPI (0.2 μg/ml) for 30 min. Cell images were captured using a Nikon. TI-E inverted fluorescence Microscope.

### Statistical analysis

All functional assays were performed at least three times independently. Data were shown as means ± SD. All statistical data were calculated using GraphPad Prism version 8.0. One-way ANOVA followed by Dunnett’s test were used to identify statistically significant differences between treatments. Significance of mean comparison is annotated as follow: ns, not significant; **, P<0.01; ***, P<0.001. A p value of < 0.05 was considered to be statistically significant.

## Supporting information

all supplementary material

## Data availability

Coordinate for the atomic structure has been deposited in the RCSB Protein Data Bank under RCSB ID XXX. The data that support the findings of this study are available from the corresponding author upon reasonable request.

## Acknowledgements

We thank Jiawei Wang for providing the suggestion for structure determination, Casey Flower and Jorge Galan for constructive proofreading of this manuscript, the staffs from BL17U1/BL18U/BL19U1 beamlines of National Facility for Protein Science Shanghai (NFPS) at Shanghai Synchrotron Radiation Facility (SSRF) for assistance during data collection, and Xiaoju Li from Shandong University Core facilities for life and environmental sciences for her help with the XRD. This work was supported by the National Natural Science Foundation of China (31901943 and 31770143), the Major Basic Program of Natural Science Foundation of Shandong Province (ZR2019ZD21), China Postdoctoral Science Foundation funded project (2019T120585 and 2019M652370), Youth Interdiscipline Innovative Research Group of Shandong University (2020QNQT009) and Taishan Young Scholars (tsqn20161005).

