## Supplementary material for "Structural insights into the insecticidal Vip3A toxin of *Bacillus thuringiensis*": all supplementary material

#### Supplementary Figures:

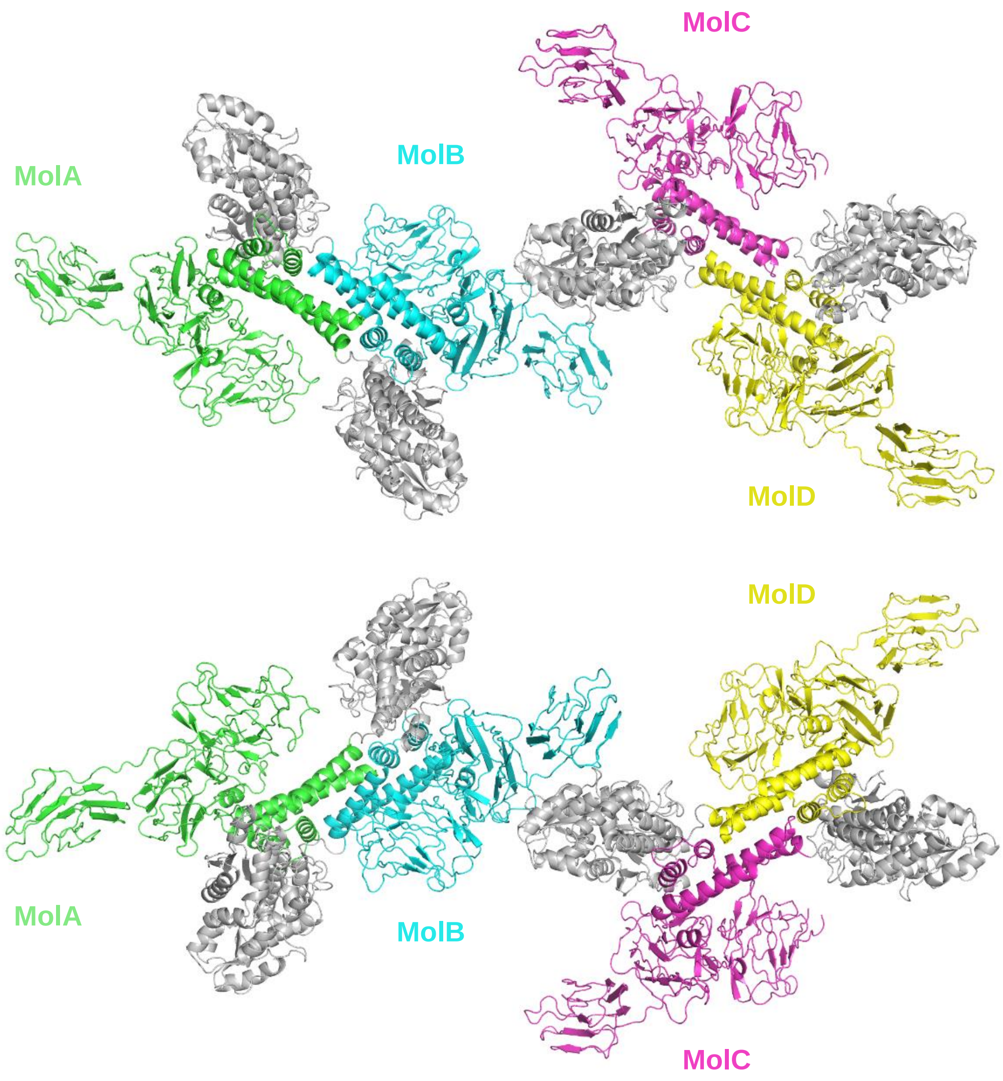


**Supplementary Figure 1. Structure of MBP-Vip3Aa11$_{200\text{-end}}$ in the *P*2$_1$ space group.** Two views of MBP-Vip3Aa11$_{200\text{-end}}$ structure in one asymmetric unit. There are four molecules of MBP-Vip3Aa11$_{200\text{-end}}$ in one asymmetric unit and they are arranged into two copies of dimer in the different orientations. The molecule A, B, C and D are shown in green, cyan, magenta and yellow, respectively. The MBP (Maltose Bind Protein) tags are shown in silver color in all four molecules. The interaction area between molecule B and C is less than 500 Å$^2$, as calculated by PISA server.

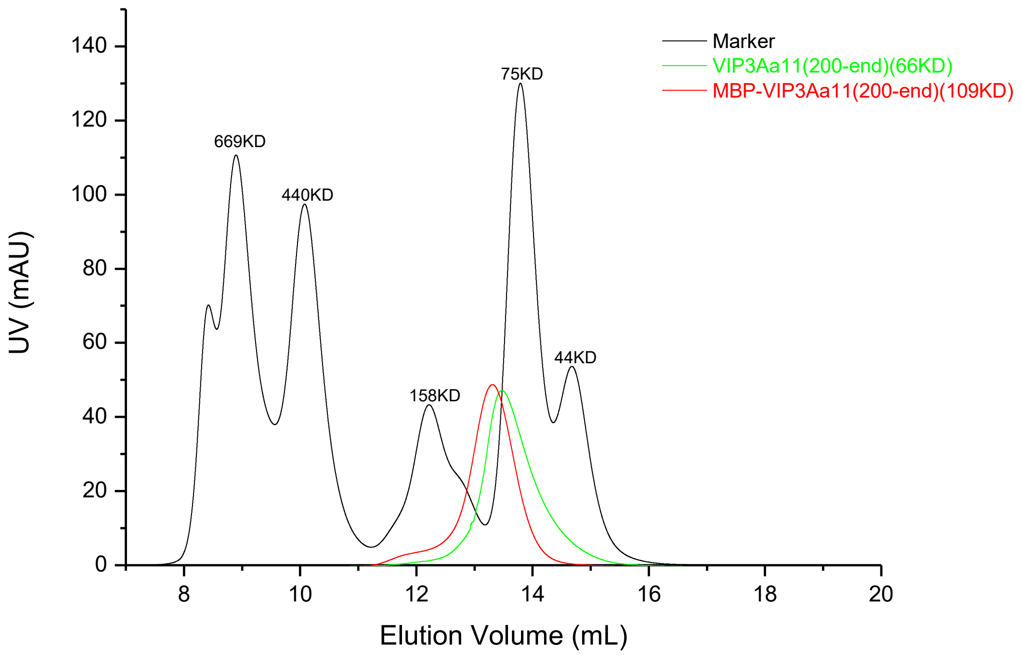


**Supplementary Figure 2. Vip3Aa11$_{200\text{-end}}$ is a monomer in solution.** Gel filtration profiles of the Vip3Aa11$_{200\text{-end}}$ protein, MBP-Vip3Aa11$_{200\text{-end}}$ protein and the molecular weight markers on Superdex-200 increase column (GE Healthcare) are shown. The sizes of the molecular weight markers are labelled on top of the peaks.

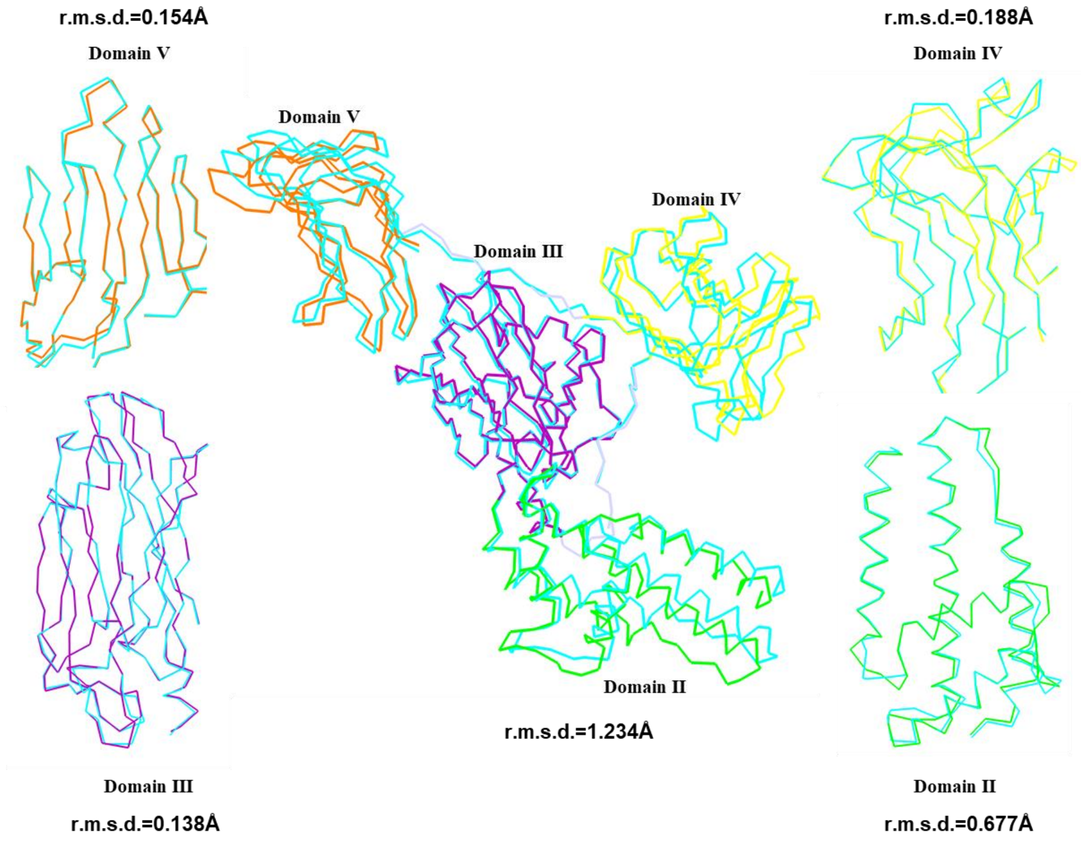


**Supplementary Figure 3. Structural alignment between Molecule A and B from the Vip3Aa11$_{200\text{-end}}$ dimer.** Structure superimposition for the Vip3Aa11$_{200\text{-end}}$ and each domain between molecule A and B from the Vip3Aa11$_{200\text{-end}}$ dimer structure. Molecule A is coloured as Figure 1A, and Molecule B is shown in cyan color. The root mean square deviation (r.m.s.d) of each alignment is listed.

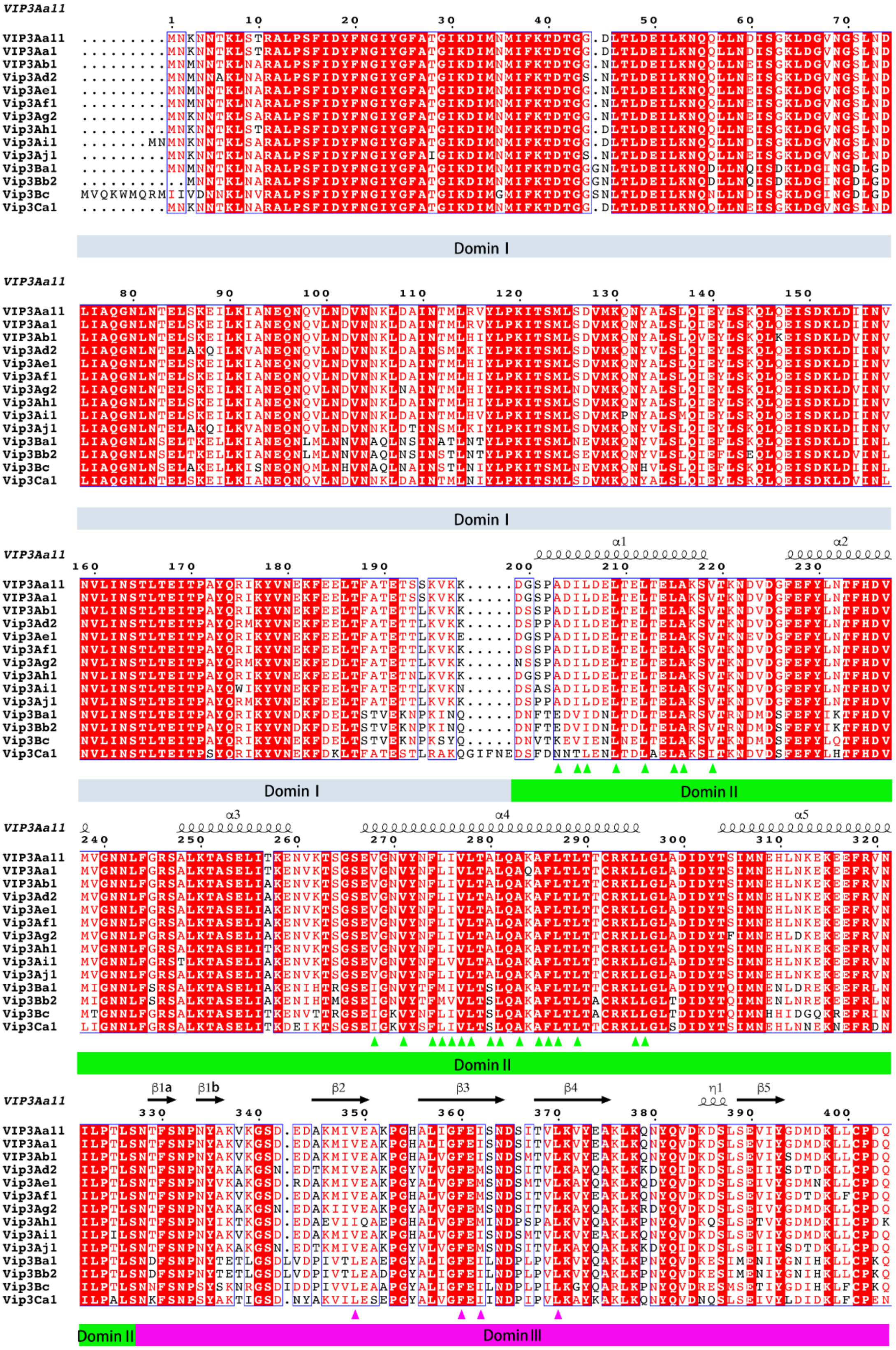
VIP3Aa11
VIP3Aa1
VIP3Ab1
Vip3Ad2
Vip3Ae1
Vip3Af1
Vip3Ag2
Vip3Ah1
Vip3Ai1
Vip3Aj1
Vip3Ba1
Vip3Bb2
Vip3Bc
Vip3Ca1
Domin I
α1
α2
Domin II
α3
α4
α5
β1a
β1b
β2
β3
β4
η1
β5
Domin III

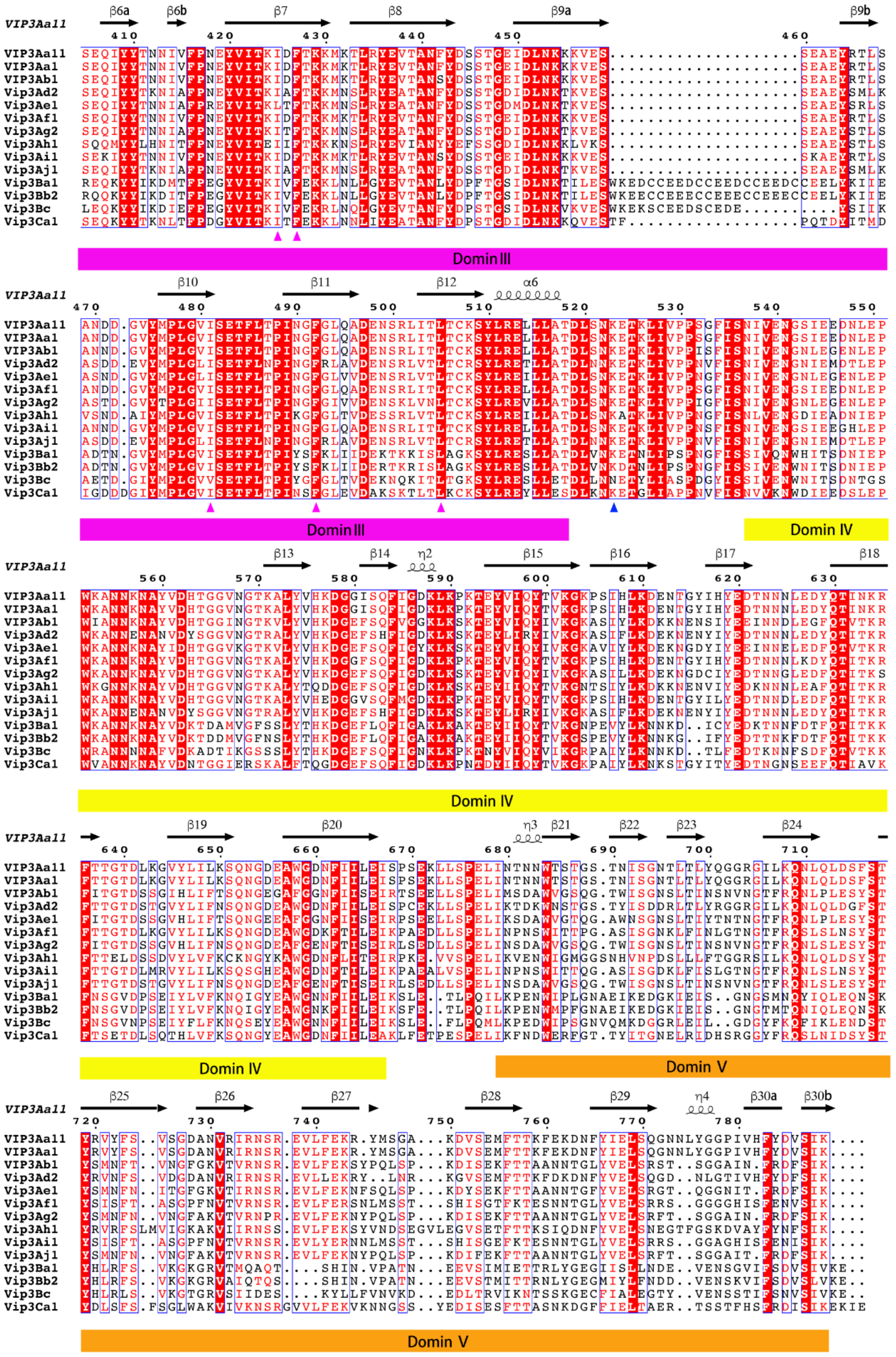

**Supplementary Figure 4. Sequence alignment of selected Vip3 family members.** Each domain is indicated by the lines bellowing sequences, coloured as in Figure A. Secondary structural elements of Vip3Aa11 are shown above the sequences. The conserved hydrophobic amino acid residues discussed in domain II and domain III are marked with green and magenta triangles, respectively. The potential cleavage site between domain III and domain IV is highlighted with blue triangle. ClustalX2 was used for the sequence alignment. ESPript-3.0 was used to generate the figure.

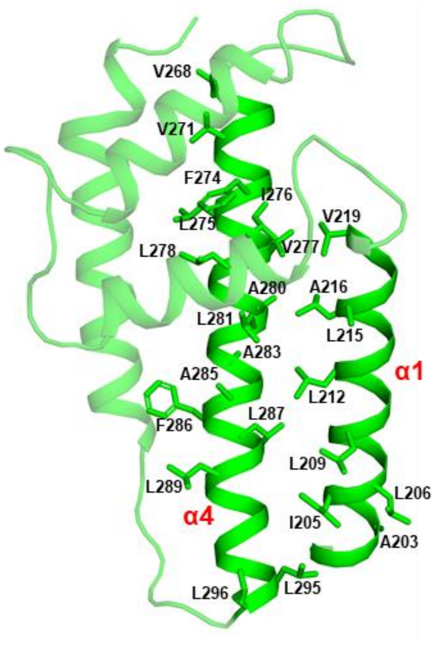


**Supplementary Figure 5. Two hydrophobic helices from domain II.** The hydrophobic amino acid residues are shown as stick and labelled with residue numbers.

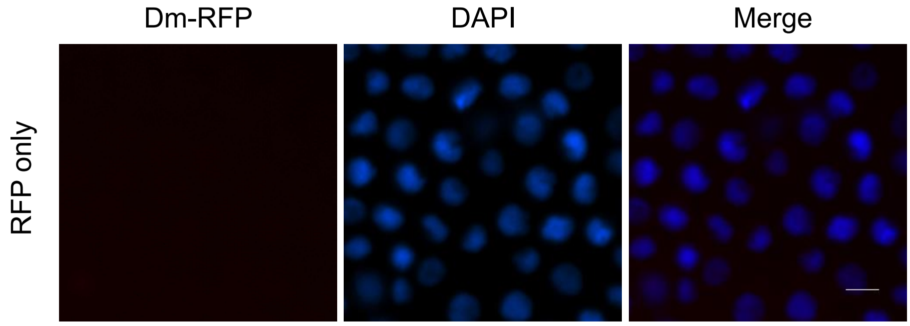


**Supplementary Figure 6. Molecule surface model of domain III of Vip3Aa11$_{200\text{-end}}$.** Fluorescence microscope images of Sf9 cells treated with RFP protein only for 6 h as control. The images are representative of three independent experiments. Nuclei are stained with DAPI (blue), Scale bar, 10 μm.

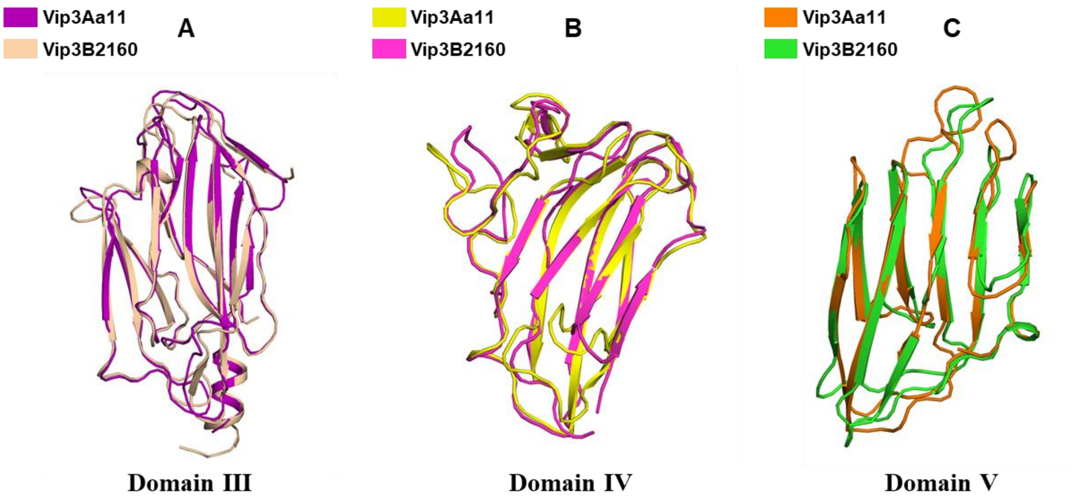


**Supplementary Figure 7. Structural comparation between corresponding domains of Vip3Aa11$_{200\text{-end}}$ and Vip3B2160.** Structure superimposition for domain III (A), domain IV (B) and domain V (C) between Vip3Aa11$_{200\text{-end}}$ and Vip3B2160. Each domain is color-coded as the indication.

Supplementary Table 1. X-ray and refinement statistics.

| | Se-SAD | Native I | Native II |
|---|---|---|---|
| **Data collection** | | | |
| Wavelength (Å) | 0.9792 | 0.9793 | 0.9793 |
| Space group | $P2_12_12_1$ | $P2_12_12_1$ | $P2_1$ |
| Cell dimensions | | | |
| *a*, *b*, *c* (Å) | 125.705,139.351,163.205 | 126.079,140.202,167.636 | 136.088,127.884,149.178 |
| α, β, γ (°) | 90, 90, 90 | 90, 90, 90 | 90,91.378,90 |
| $R_{meas}$ | 0.153 (0.829) | 0.207(0.915) | 0.145 (0.753) |
| Mean $I/\sigma I$ | 9.30(2.80) | 7.47(1.32) | 6.11 (1.00) |
| CC1/2 | 0.981(0.674) | 0.983(0.565) | 0.993(0.541) |
| Completeness (%) | 91.25 (79.60) | 93.28(91.40) | 97.33 (95.63) |
| Redundancy | 3.5 (3.7) | 3.9(3.8) | 2.3 (2.3) |
| Wilson B-factor | 48.45 | 99.23 | 71.43 |
| **MR-SAD Phasing** | | | |
| *Selenium sites (ShelX/D)* | 11 | | |
| PATFOM (/D) | 24.47 | | |
| Overall CC (ShelX/E) (%) | 6.57 | | |
| Pseudo-Free CC (/E) (%) | 60.29 | | |
| Final Map CC (/E) | 0.995(0.897) | | |
| Figure of Merit (FOM) | 0.564 | | |
| **Refinement** | | | |
| Resolution (Å) | 49.93 - 3.91 (4.04-3.90) | 42.42 -3.62 (3.75 - 3.62) | 28.93 - 3.195 (3.309 - 3.195) |
| No. reflections | | | 82266 |
| $R_{work}$ / $R_{free}$ | | | 0.1980/0.2389 |
| No. atoms | | | 30179 |
| Protein | | | 29960 |
| Ligand/ion | | | 42 |
| Water | | | 177 |
| Average *B*-factors | | | 74.14 |
| Protein | | | 74.32 |
| Ligand | | | 86.70 |
| Water | | | 40.48 |
| R.m.s. deviations | | | |
| Bond lengths (Å) | | | 0.002 |
| Bond angles (°) | | | 0.58 |
| Ramachandran | | | |
| Favored (%) | | | 95.01 |
| Allowed (%) | | | 4.80 |
| Outliers (%) | | | 0.18 |

Number of crystals for each structure should be noted in footnote.

[a] Values in parentheses are for highest-resolution shell.

### Supplementary Table 2. Glycan array screening results

| Glycan ID | Glycan Structures | Glycan Name | Ave (n=3) | StdDev (n=3) |
|---|---|---|---|---|
| 1 | D-Mannose | D-Mannose | 3473.66 | 1130.21 |
| 2 | D-Glucose | D-Glucose | 3373.5 | 310.78 |
| 3 | D-Fucose | D-Fucose | 12426.33 | 1895.9 |
| 4 | D-Galactose | D-Galactose | 7309 | 3438.55 |
| 5 | L-Fucose | L-Fucose | 13209 | 414.89 |
| 6 | L-Rhamnose | L-Rhamnose | 15706.33 | 1194.84 |
| 7 | D-ManNAc | D-ManNAc | 5921 | 2682.55 |
| 8 | D-GlcNAc | D-GlcNAc | 7525 | 3576.12 |
| 9 | Neu5Ac | Sialic acid | 113.5 | 29.04 |
| 10 | Galβ1-4(Fucα1-3)Glc | 3FL | 651.33 | 151.58 |
| 11 | Galβ1-4Glc | Lactose | 671.83 | 94.28 |
| 12 | Galβ1-4GlcNAc | LacNAc | 719.16 | 93.97 |
| 13 | Galβ1-3GlcNAc | Lac-N-biose | 1068.66 | 892.64 |
| 14 | Galβ1-4Gal | 4-β-Galactobiose | 1210 | 866.3 |
| 15 | GalNAcβ1-3Gal | β-D-N-acetyl-galactosaminyl 1-3 galactose | 374.33 | 185.43 |
| 16 | Glcα1-4Glc | Maltose | 1349.66 | 1627.49 |
| 17 | Glcβ1-4Glc | Cellobiose | 771.66 | 172.24 |
| 18 | GlcNAcβ1-4GlcNAc | Diacetylchitobiose | 326.33 | 42.53 |
| 19 | GlcNAcβ1-2Man | β-D-N-acetylglucosaminyl 1-2 mannose | 227.16 | 20.32 |
| 20 | GlcNAcβ1-4MurNAc | GlcNAcMurNAc | 184.5 | 33.23 |
| 21 | GlcNH$_2$β1-4GlcNH$_2$ | Chitobiose | 4516.33 | 1609.2 |
| 22 | Manα1-2Man | 2-Mannbiose | 544 | 280.44 |
| 23 | Manα1-3Man | 3-Mannobiose | 1041.16 | 636.74 |
| 24 | Manα1-4Man | 4-Mannobiose | 977.83 | 1126.39 |
| 25 | Manα1-6Man | 6-Mannobiose | 98.66 | 38.59 |
| 26 | Fucα1-2Gal | Blood H Disaccharide | 98.33 | 122.46 |
| 27 | ΔGlcA(2S)α1-4GlcNS(6S) | Heparin Disaccharide | 47.5 | 68.51 |
| 28 | GlcNH$_2$β1-4GlcNH$_2$β1-4GlcNH2 | Chitotriose | 5943.16 | 865.18 |
| 29 | Galβ1-4Galβ1-4Glc | Globotriose (P$^K$ antigen) | 190 | 58.07 |
| 30 | GlcNAcβ1-4GlcNAcβ1-4GlcNAc | Triacetyl chitotriose | 4194.5 | 1002 |
| 31 | Neu5Acα2-3Galβ1-4Glc | 3'-SL (GM3 Glycan) | 153.16 | 28.55 |
| 32 | Neu5Acα2-6Galβ1-4Glc | 6'-SL | 147.16 | 63.17 |
| 33 | Neu5Acα2-3Galβ1-4GlcNAc | 3'-SLN | 236.33 | 15.59 |
| 34 | Neu5Acα2-6Galβ1-4GlcNAc | 6'-SLN | 207.66 | 50.65 |
| 35 | SO$_3$-3Galβ1-4(Fucα1-3)GlcNAc | Sulpho-Lewis x | 149.66 | 122.32 |
| 36 | SO$_3$-3Galβ1-3(Fucα1-4)GlcNAc | Sulpho-Lewis a | 18.33 | 74.65 |
| 37 | Galβ1-4(Fucα1-3)GlcNAc | Lewis x Trisaccharide | 531.5 | 121.26 |
| 38 | Galβ1-3(Fucα1-4)GlcNAc | Lewis a Trisaccharide | 236.66 | 174.61 |
| 39 | Glcα1-4Glcα1-4Glc | Maltotriose | 110.33 | 94.89 |
| 40 | Glcβ1-4Glcβ1-4Glc | Cellotriose | 110.66 | 41.93 |
| 41 | Fucα1-2Galβ1-4Glc | 2'FL | 45.66 | 14.29 |
| 42 | Galβ1-3Galβ1-3GlcNAc | Linear B-2 Tri (Blood Group B Type 2 Linear Tri) | 132.33 | 95.25 |
| 43 | Galα1-4Galβ1-4GlcNAc | P1 antigen Tri | 27.66 | 20.79 |
| 44 | Fucα1-2Galβ1-3GlcNAc | Blood group H Trisaccharide | 15.66 | 28.36 |
| 45 | GalNAcα1-3-(Fucα1-2)Gal | Blood group A Trisaccharide | -20 | 30.02 |
| 46 | Galα1-3(Fucα1-2)Gal | Blood group B Trisaccharide | 288 | 353.28 |
| 47 | Galβ1-3GlcNAcβ1-3Galβ1-4Glc | Lacto-N-tetraose (LNT) | 6303.33 | 606.74 |
| 48 | Galβ1-4GlcNAcβ1-3Galβ1-4Glc | Lacto-N-neotetraose (LNnT) | 81 | 11.37 |
| 49 | Galα1-3Galβ1-4Galβ1-3Gal | Gal4 | 214.66 | 67.11 |
| 50 | Glcα1-4Glcα1-4Glcα1-4Glc | Maltotetratose | 161 | 52.39 |
| 51 | Neu5Acα2-3Galβ1-4(Fucα1-3)GlcNAc | Sialyl Lewis x | 42.66 | 41.02 |
| 52 | Neu5Acα2-3Galβ1-3(Fucα1-4)-GlcNAc | Sialyl Lewis a | 28 | 16.17 |
| 53 | Fucα1-2Galβ1-3(Fucα1-4)GlcNAc | Lewis b Tetrasaccharide | 41.33 | 10.15 |
| 54 | Fucα1-2Galβ1-4(Fucα1-3)GlcNAc | Lewis y Tetrasaccharide | 26.33 | 19.05 |
| 55 | Fucα1-2Galβ1-3GlcNAcβ1-3Galβ1-4Glc | Lacto-N-Fucopentaose I (LNFP-I) | 20.33 | 24.27 |
| 56 | Galβ1-3(Fucα1-4)GlcNAcβ1-3Galβ1-4Glc | Lacto-N-Fucopentaose II (LNFP-II) | 37 | 32.5 |
| 57 | Galβ1-4(Fucα1-3)GlcNAcβ1-3Galβ1-4Glc | Lacto-N-Fucopentaose III (LNFP-III) | 115.66 | 52.6 |
| 58 | Fucα1-2Galβ1-4(Fucα1-3)GlcNAcβ1-3Gal | Lewis Y Pentasaccharide | 92 | 23.46 |
| 59 | Glcα1-4Glcα1-4Glcα1-4Glcα1-4Glc | Maltopentaose | 66.66 | 20.21 |
| 60 | [Manα1-3-(Manα1-6)-Manα1-6]- (Manα1-3)-Man | Man5 | 18.66 | 13.65 |
| 61 | Neu5Aca2-6(Galβ1-3)GlcNAcβ1-3Galβ1-4Glc | LS-Tetrasaccharide b (LsTb) | -3.67 | 8.54 |
| 62 | Neu5Aca2-6Galβ1-4GlcNAcβ1-3Galβ1-4Glc | LS-Tetrasaccharide c (LsTc) | 3.33 | 22.65 |
| 63 | Galβ1-3GalNAcβ1-4(Neu5Acα2-3)Galβ1-4Glc | GM1 Glycan | -21 | 29.74 |
| 64 | GalNAcα1-3(Fucα1-2)Galβ1-4(Fucα1-3)Glc | Blood Group A Pentasaccharide | 236 | 195.47 |
| 65 | (GlcNAcβ1-2Manα1)$_2$-3,6Man | Bianntennary N-linked Core Pentasaccharide | 53 | 43.02 |
| 66 | Galβ1-3(Fucα1-4)GlcNAcβ1-3Galβ1-4(Fucα1-3)Glc | Lacto-N-difucohexaose II (LNDFH II) | 99.66 | 32.65 |
| 67 | Neu5Aca2-3Galβ1-3(Neu5Aca2-6)GlcNAcβ1-3Galβ1-4Glc | DSLNT | 69.66 | 69.66 |
| 68 | GlcNAcβ1-4GlcNAcβ1-4GlcNAcβ1-4GlcNAcβ1-4GlcNAcβ1-4GlcNAc | Hexaacetyl Chitohexaose | 21 | 46.69 |
| 69 | Glcα1-4Glcα1-4Glcα1-4Glcα1-4Glcα1-4Glc | Maltohexaose | 38.66 | 21.78 |
| 70 | Glcα1-4Glcα1-4Glcα1-4Glcα1-4Glcα1-4Glcα1-4Glc | Maltoheptaose | 67.66 | 127.03 |
| 71 | (4GlcAβ1-4GlcNAc(6S)α1)$_4$ | Heparin Octasaccharide | 16.66 | 16.92 |
| 72 | Manα1-6(Manα1-3)Manα1-6(Manα1-3)Manβ1-4GlcNAcβ1-4GlcNAc | MAN-5; (Man)5(GlcNAc)2 | 6.33 | 18.52 |
| 73 | Manα1-2Manα1-6(Manα1-2Manα1-3)Manα1-6(Manα1-2Manα1-2Manα1-3)Manβ1-4GlcNAcβ1-4GlcNAc | MAN-9; (Man)9(GlcNAc)2 | -0.67 | 10 |
| 74 | GalNAcα1-3(Fucα1-2)Galβ1-3GlcNAc | Blood Group A Type 1 Tetrasaccharide | -5.34 | 5.03 |
| 75 | GalNAcα1-3(Fucα1-2)Galβ1-4GlcNAc | Blood Group A Type 2 Tetrasaccharide | 20.66 | 12.9 |
| 76 | GalNAcα1-3(Fucα1-2)Galβ1-3GalNAc | Blood Group A Type 3/4 Tetrasaccharide | 77 | 132.61 |
| 77 | Galα1-3-(Fuc1-2)Galβ1-4(Fucα1-3)Glc | Blood Group B Pentasaccharide | -14 | 20.26 |
| 78 | Galα1-3-(Fuc1-2)Galβ1-3GlcNAc | Blood Group B Type 1 Tetrasaccharide | 10.66 | 50.54 |
| 79 | Galα1-3-(Fuc1-2)Galβ1-4GlcNAc | Blood Group B Type 2 Tetrasaccharide | -28.67 | 17.06 |
| 80 | Galα1-3-(Fuc1-2)Galβ1-3GalNAc | Blood Group B Type 3/4 Tetrasaccharide | -11 | 15.04 |
| 81 | Neu5Acα2-3Galβ1-4(Fucα1-3)GlcNAcβ1-3Gal | Sialyl Lewis x Pentasaccharide | -28.67 | 11.36 |
| 82 | Neu5Acα2-3(GalNAcβ1-4)Galβ1-4Glc | GM2 Glycan | 13 | 13.58 |
| 83 | Neu5Ac2-3Galβ1-3GlcNAcβ1-3Galβ1-4Glc | LS-Tetrasaccharide a (LsTa) | 5.33 | 10.58 |
| 84 | Neu5Acα2-3Galβ1-4GlcNAcβ1-3Galβ1-4Glc | LS-Tetrasaccharide d (LsTd) | 13.66 | 23.03 |
| 85 | Neu5Acα2-3Galβ1-3GalNAcβ1-3Galα1-4Galβ1-4Glc | Stage-specific Embryonic Antigen 4 (SSEA-4) | 56 | 69.5 |
| 86 | Glcβ1-4Glcβ1-4Glcβ1-4Glc | Cellotetraose | -9 | 10.02 |
| 87 | Glcβ1-4Glcβ1-4Glcβ1-4Glcβ1-4Glc | Cellopentaose | -20.67 | 19.67 |
| 88 | Glcβ1-4Glcβ1-4Glcβ1-4Glcβ1-4Glcβ1-4Glc | Cellohexaose | -37.34 | 7.51 |
| 89 | D-GalNAc | D-GalNAc | -3 | 40.77 |
| 90 | Galβ1-3GalNAc | T antigen | -22.67 | 11.53 |
| 91 | GalNAcα1-3Gal | Adi | 11.33 | 14.11 |
| 92 | GalNAcα1-3Galβ1-4Glc | α-D-N-Acetylgalactosaminyl 1-3 galactose β 1-4 glucos | -3.67 | 4.58 |
| 93 | Galβ1-6Gal | β 1-6 galactobiose | 30 | 39.58 |
| 94 | GalNAcα1-3GalNAc | Forssman disaccharide | -9.34 | 15.63 |
| 95 | GalNAcβ1-4Gal | Receptor for pili of Pseudomonas aeruginosa | 52.33 | 70.49 |
| 96 | Galβ1-3GalNAcβ1-3Galα1-4Galβ1-4Glc | Stage-specific Embryonic Antigen 3 (SSEA-3) | 0.66 | 41.48 |
| 97 | Fucα1-2Galβ1-3GalNAcβ1-3Galα1-4Galβ1-4Glc | Globo-H | 2.33 | 70.49 |
| 98 | Manα1-6(Manα1-3)Manα1-6(Manα1-2Manα1-3)Manβ1-4GlcNAcβ1-4GlcNAc | MAN-6; (Man)6(GlcNAc)2 | 17.66 | 99.36 |
| 99 | [Manα1-2]Manα1-6(Manα1-3)Manα1-6(Manα1-2Manα1-3)Manβ1-4GlcNAcβ1-4GlcNAc | MAN-7; (Man)7(GlcNAc)2 | -43 | 24.79 |
| 100 | [Manα1-2] [Manα1-2] Manα1-6(Manα1-3)Manα1-6(Manα1-2Manα1-3)Manβ1-4GlcNAcβ1-4GlcNAc | MAN-8; (Man)8(GlcNAc)2 | 25.66 | 45.54 |

Supplementary Table 3. Primers used in this study

| Primer name | Sequence(5′→3′) | Function |
|---|---|---|
| Vip200-F | GAGCTTTCGCTGCAGCGTCCGGCTCTCCTGCAGATATTC | Vip$_{200\text{-}end}$ cloning |
| Vip200-R | GTTAGCAGCCGGATCTCAGTGTTACTTAATAGAGACATCGTAAAAATGTAC | Vip$_{200\text{-}end}$ cloning |
| DmI-III-F | AAGAAGGAGATATACCATGGGCATGAACAAGAATAATACTAAATTAAGC | DmI-III cloning |
| DmI-III-R | TCGACTGCAGAGGCCTGCATAGAAAGTGTAGGGAGGATGTTTAC | DmI-III cloning |
| DmVI-V-F | AAGAAGGAGATATACCATGGGCGGTTTTATTAGCAATATTGTAGAG | DmVI-V cloning |
| DmVI-V-R | TCGACTGCAGAGGCCTGCATCTTAATAGAGACATCGTAAAAATG | DmVI-V cloning |
| DmI-II-F | AAGAAGGAGATATACCATGGGCATGAACAAGAATAATACTAAATTAAGC | DmI-II cloning |
| DmI-II-R | TCGACTGCAGAGGCCTGCATAGAAAGTGTAGGGAGGATGTTTAC | DmI-II cloning |
| DmII-III-F | AAGAAGGAGATATACCATGGGCGATGGCTCTCCTGCAGATATTCTTG | DmII-III cloning |
| DmII-III-R | TCGACTGCAGAGGCCTGCATAGAAAGTGTAGGGAGGATGTTTAC | DmII-III cloning |
| DmIII-F | AAGAAGGAGATATACCATGGGCACACTTTCTAATACTTTTTCTAATC | DmIII cloning |
| DmIII-R | TCGACTGCAGAGGCCTGCATTTCTTTATTGCTTAAGTCTGTTGC | DmIII cloning |
| pET-MBP-F | CACTGAGATCCGGCTGCTAAC | pET28-MBP cloning |
| pET-MBP-R | TTCTTTATTGCTTAAGTCTG | pET28-MBP cloning |
| pET-RFP-F | ATGCAGGCCTCTGCAGTCGACGGG | pET28-RFP cloning |
| pET-RFP-R | GCCCATGGTATATCTCCTTCTT | pET28-RFP cloning |

The nucleic acid bases corresponding to each gene are labeled with underscores.
